# Fast and accurate single-cell RNA-Seq analysis by clustering of transcript-compatibility counts

**DOI:** 10.1101/036863

**Authors:** Vasilis Ntranos, Govinda M. Kamath, Jesse Zhang, Lior Pachter, David N. Tse

## Abstract

Current approaches to single-cell transcriptomic analysis are computationally intensive and require assay-specific modeling which limit their scope and generality. We propose a novel method that departs from standard analysis pipelines, comparing and clustering cells based not on their transcript or gene quantifications but on their transcript-compatibility read counts. In re-analysis of two landmark yet disparate single-cell RNA-Seq datasets, we show that our method is up to two orders of magnitude faster than previous approaches, provides accurate and in some cases improved results, and is directly applicable to data from a wide variety of assays.

## Introduction

Single-cell RNA-Seq (scRNA-Seq) has proved to be a powerful tool for probing cell states [1-5], defining cell types [6-9], and describing cell lineages [10-13]. These applications of scRNA-Seq all rely on two computational steps: quantification of gene or transcript abundances in each cell and clustering of the data in the resulting abundance x cell expression matrix [14,15]. There are a number of challenges in both of these steps that are specific to scRNA-Seq analysis. While methods for transcript/gene abundance estimation from bulk RNA-Seq have been extensively tested and benchmarked [16], the wide variety of assay types in scRNA-Seq [17-25] have required a plethora of customized solutions [2,6,7,9,11-13,24,26-37] that are difficult to compare to each other. Furthermore, the quantification methods used all rely on read alignment to transcriptomes or genomes, a time consuming step that will not scale well with the increasing numbers of reads predicted for scRNA-Seq [15,38]. Clustering based on scRNA-Seq expression matrices can also require domain specific information, e.g. temporal information [33] or functional constraints [37] so that in some cases hand curation of clusters is performed after unsupervised clustering [7].

In [39], a method of collapsing bulk read alignments into “equivalence classes” of reads was introduced for the purpose of estimating alternative splicing isoform frequencies from bulk RNA-Seq data. Each equivalence class consists of all the reads that are compatible with the same set of transcripts. (See Figure 1 for an example.) The collapsing of reads into equivalence classes was initially introduced to allow for significant speedup of the E-step in the expectation-maximization (EM) algorithm used in some RNA-Seq quantification programs [40,41], as the read counts in the equivalence classes, or *transcript-compatibilty counts* (TCC), correspond to the sufficient statistics for a standard RNA-Seq model [42]. In other words, the use of transcript-compatibility counts was an intermediate computation step towards quantifying transcript abundances. In this paper we instead consider the direct use of such counts for the comparison and clustering of scRNA-Seq cells. Figure 2 shows an outline of a method we have developed for clustering and analyzing scRNA-Seq data; the key idea is to base clustering not on the quantification of transcripts or genes but on the transcript-compatibility counts for each cell. We note that equivalence classes have also been used in [43, 44] to define similarity scores between de novo assembled transcripts.

**Figure 1.**
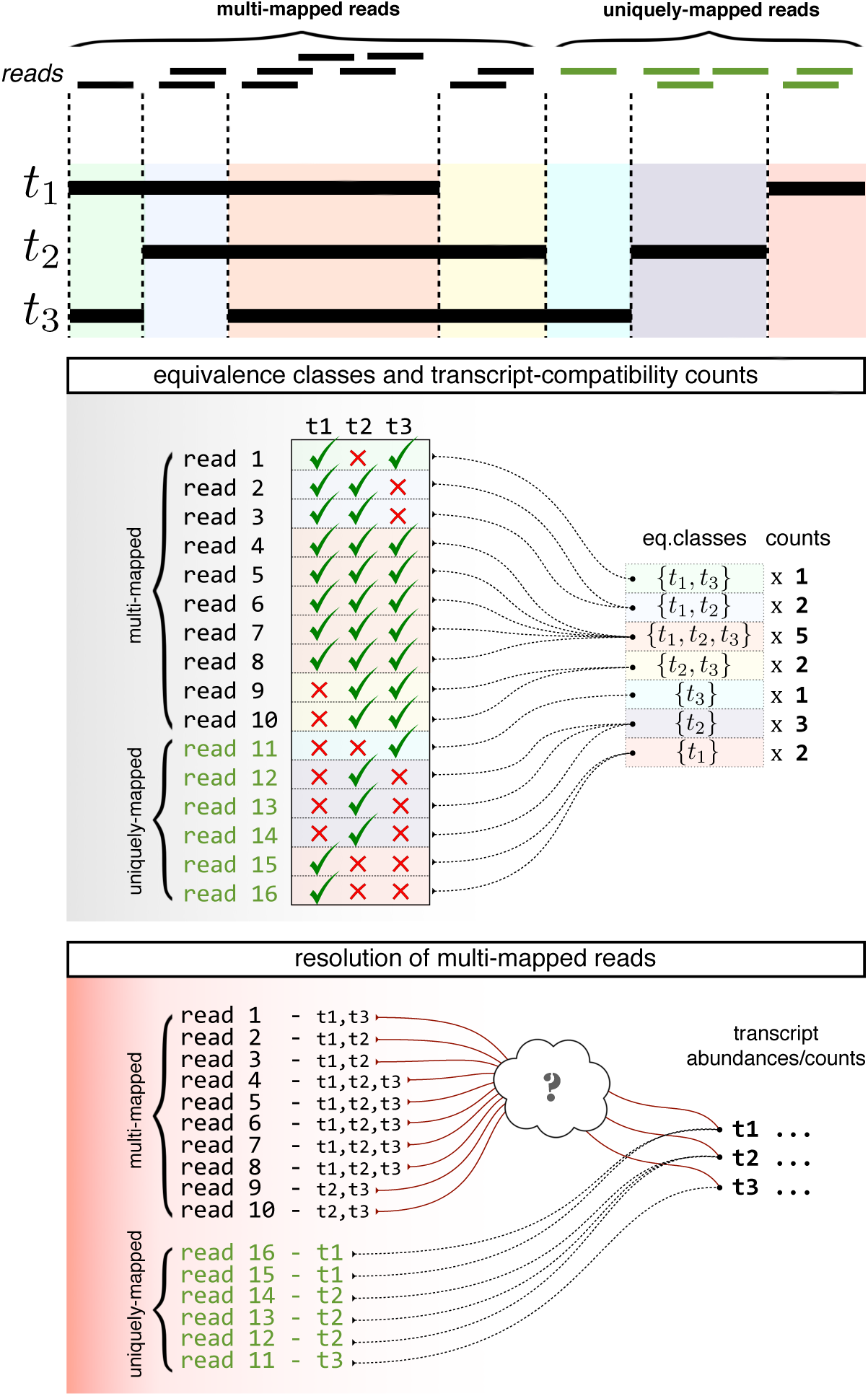
Equivalence class and transcript-compatibility counts. This figure gives an example of how reads are collapsed into equivalence classes. Each read is mapped to one or more transcripts in the reference transcriptome; these are transcripts that the read is compatible with, i.e. the transcripts that the read could possibly have come from. For example, read 1 is compatible with transcripts t1 and t3, read 2 is compatible with transcripts t1 and t2, and so on. An equivalence class is a group of reads that is compatible with the same set of transcripts. For example, reads 4,5,6,7,8 are all compatible with t1,t2 and t3 and they form an equivalence class. Since the reads in an equivalence class are all compatible with the same set of transcripts, we simply represent an equivalence class by that set of transcripts. For example, the equivalence class consisting of reads 4,5,6,7,8 is represented by {*t*1, *t*2, *t*3}. Aggregating the number of reads in each equivalence class yields the corresponding transcript-compatibility counts. Note that in order to estimate the transcript abundances from the transcript-compatibility counts, a read-generation model is needed to resolve the multi-mapped reads.

**Figure 2.**
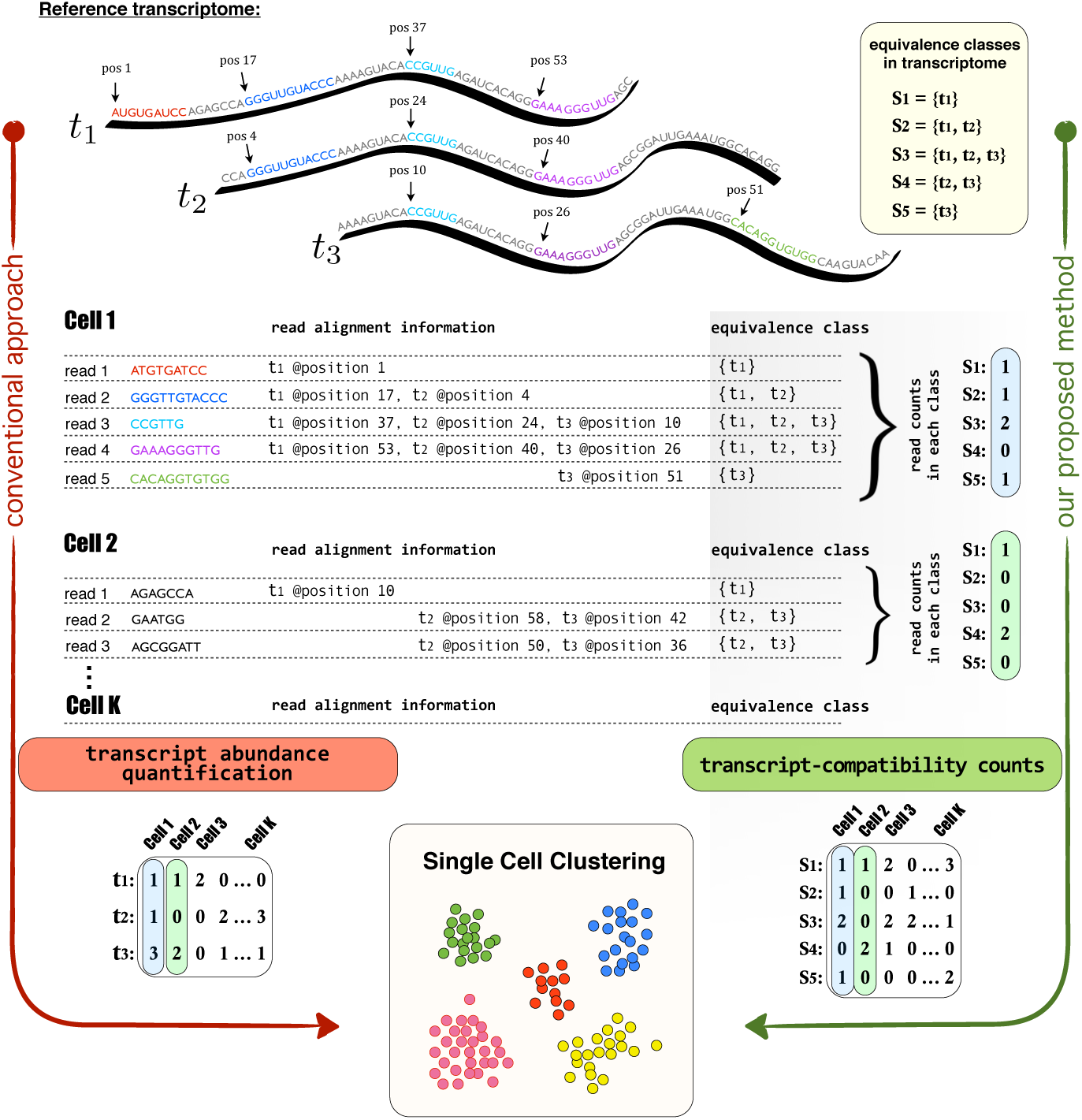
Overview of the Method. This figure illustrates our transcript-compatibility counts (abbreviated TCC) clustering method in a very simple, yet instructive example and highlights its major differences with respect to the conventional single cell clustering approach. Here, we consider an scRNA-Seq example with K cells (only the reads coming from Cell1 and Cell2 are shown here) and a reference transcriptome consisting of three transcripts, t_1_, t_2_ and t_3_. Conventional approach: Single cells are clustered based on their transcript or gene abundances (here we only focus on transcripts for concreteness). This widely adopted pipeline involves computing a (#transcripts × #cells) expression matrix by first aligning each cell’s reads to the reference. The corresponding alignment information is next to each read, which for the purpose of illustration only contains the mapped positions (the aligned reads of Cell1 are also annotated directly on the transcripts). While reads 1 and 5 are uniquely mapped to transcripts 1 and 3, reads 2, 3 and 4 are mapped to multiple transcripts (multi-mapped reads). The quantification step must therefore take into account a specific read-generating model and handle multi-mapped reads accordingly. Our proposed method: Single cells are clustered based on their transcript-compatibility counts. Our method assigns the reads of each cell to equivalence classes via the process of pseudoalignment and simply counts the number of reads that fall in each class to construct a (#eq.classes × #cells) matrix of transcript-compatibility counts. Then, the method proceeds by directly using the transcript-compatibility counts for downstream processing and single cell clustering. The underlying idea here is that even though equivalence classes may not have an explicit biological interpretation, their read counts can collectively provide us with a distinct signature of each cell’s gene expression; transcript-compatibility counts can be thought of as feature vectors and cells can be identified by their differential expression over these features. Compared to the conventional approach, our method does not attempt to resolve multi-mapped reads (no need for an assay-specific read-generating model) and only requires transcript compatibility information for each read (no need for exact read alignment).

To better understand the relevance of transcript-compatibility counts, consider their relationship to “gene-level” counts used in many RNA-Seq analyses. In the same way that “genes” represent groupings of transcripts [45], equivalence classes as introduced by [39] are also groups of transcripts. However while the former is a biologically motivated construction, the latter is technical, consisting of groupings that capture the extent of ambiguous multiple mappings among reads. The lack of direct biological interpretation of equivalence classes makes transcript-compatibility counts less intuitive; however, as we will show, there are two significant advantages to working with them: 1) unlike transcript or gene-level quantifications, transcript-compatibility counts can be computed without a read-generation model, and hence a single clustering pipeline based on transcript-compatibility counts can be used across a wide range of scRNA-Seq assays; 2) transcript-compatibility counts can be computed by pseudoalignment, a process that does not require read alignment and can be done extremely efficiently [41].

To demonstrate both the general applicability of our method as well as its accuracy, we re-analyzed data from two recently published scRNA-Seq papers: the pseudotemporal ordering of primary human myoblasts by [12] and the cell classification in the mouse cortex and hippocampus by [7]. We show that not only are we able to recapitulate the analyses of the papers two orders of magnitude faster than previously possible, but we also provide a refinement of the published results, suggesting that our approach is both fast and accurate. To demonstrate both the general applicability of our method as well as its accuracy, we re-analyzed data from two recently published scRNA-Seq papers: the pseudotemporal ordering of primary human myoblasts by [12] and the cell classification in the mouse cortex and hippocampus by [7]. The speedup of our method makes single-cell RNA-Seq analysis interactive for the first time: sensitivity of results to parameters and annotations can be easily explored and analyses can be easily reproduced by individuals without access to significant compute resources. Furthermore, the efficiency of our methods will take on increasing significance as single-cell RNA sequencing scales to experiments with hundreds of thousands of cells and improved technologies make the acquisition of single-cell data easier and faster (for example [46]). In addition, we also illustrate the advantages of the broad applicability of our approach via its suitability to a multitude of assays. Existing pipelines must be tailored to specific assays making it difficult to perform meta-analyses and to compare results across experiments.

## Results

### Transcript-compatibility counts from pseudoalignments

To demonstrate the effectiveness of transcript-compatibility counts for scRNA-Seq analysis, we first examined how efficiently they can be computed. While transcript-compatibility counts can be extracted from read alignments (e.g. in SAM/BAM format), they do not require the full information contained in alignments. Instead, we examined the speedup possible with pseudoalignment [41], which obtains for each read the set of transcripts it is compatible with and therefore can be directly used to obtain transcript-compatibility counts.

Figure 3 shows the speed of obtaining transcript-compatibility counts via pseudoalignment in comparison to the time required to quantify RNA-Seq data with other approaches. The key result relevant for single-cell analysis is the scalability of pseudoalignment for obtaining transcript-compatability counts (Figure 3 and Supplementary Figure 2). The fixed extra cost for aligning (rather than pseudoaligning) reads for each cell is small, but when extrapolated to hundreds of thousands of cells becomes a significant (computational) cost.

**Figure 3.**
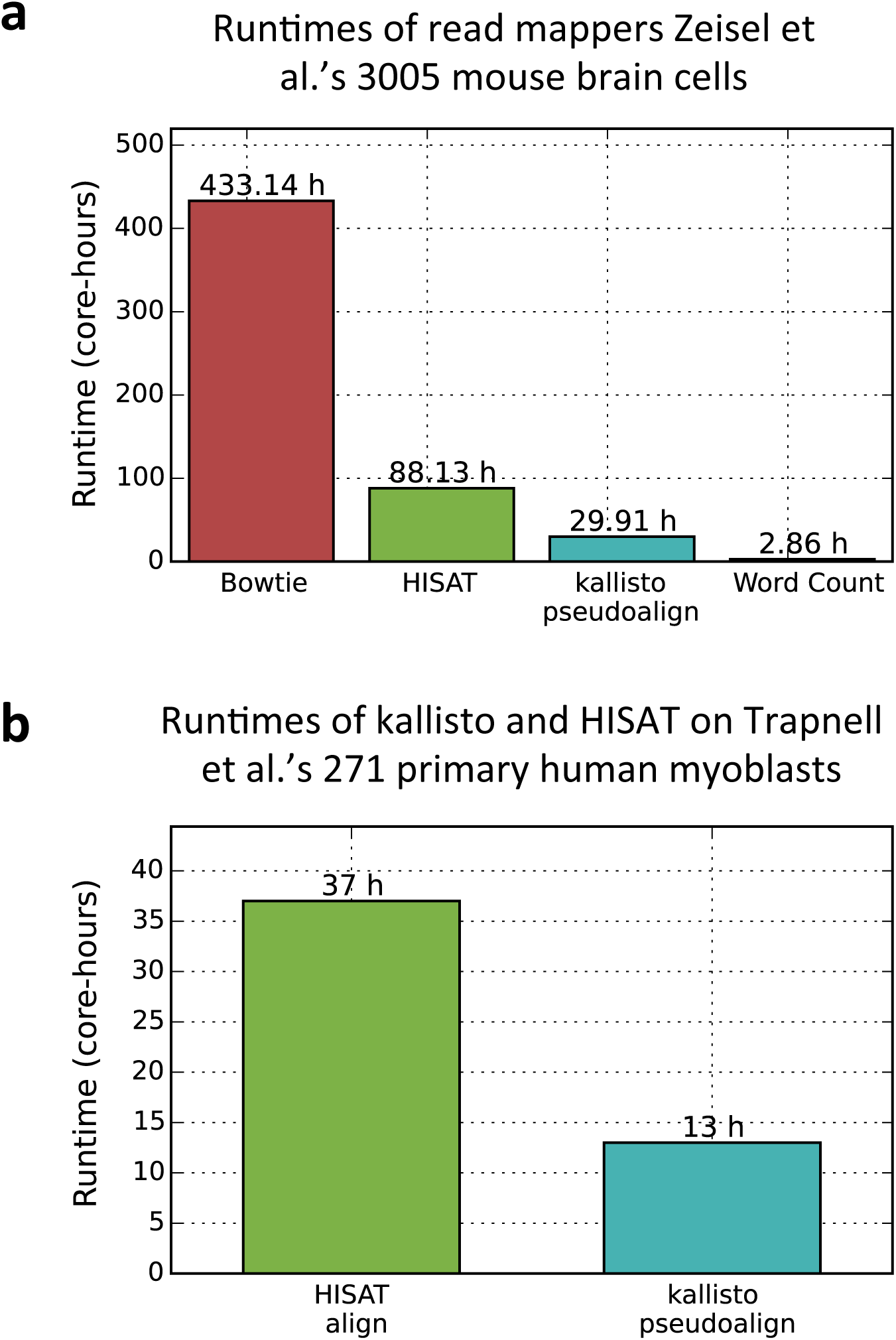
Runtime comparison of alignment methods. **(a)** The time required to process 3005 cells from mouse brain cell dataset [7] in core hours is shown here. The time taken for read alignment with Bowtie and HISAT is much larger than the time taken for kallisto pseudoalignment (which is used by our method to obtain the transcript-compatibility counts). kallisto pseudoalignment and HISAT were run on 32 cores. Bowtie and word count were each timed on 1 core with 10 randomly selected cells. The bars shown here are estimates obtained by multiplying these times by 300.5. Because we do not account for the overhead associated with parallelizing, the Bowtie and word count estimates are lower bounds on their run times in practice. After preprocessing the UMIs, each of the 5, 914, 602, 849 single-end reads in the dataset were less than 50 bp long. *(b)* The time required to process 271 cells of the dataset of [12] in core hours is shown here. As before, the time taken for read alignment with HISAT is significantly larger than the time taken for kallisto pseudoalignment. Both methods were run on 32 cores. The dataset has 814,344, 693 paired-end reads, and each mate in a pair is 100 bp long.

### Pseudotime for differentiating human myoblasts

The recently published Monocle software [12] that builds on the Cufflinks program [47] is rapidly becoming a standard tool for scRNA-Seq analysis. We therefore sought to compare our approach to Monocle, and in order to do so began with a re-analysis of the data in [12]. Figure 4 shows the temporal ordering of differentiating primary human myoblasts using transcript-compatibility counts clustering based on the Jensen-Shannon metric and the affinity propagation algorithm (see Methods). We note that unlike Cufflinks, which consists of an explicit model of RNA-Seq suitable for the data in [12] but not necessarily for other assays, our transcript-compatibility counts make no assumption about the nature of the data. Furthermore, while the re-analysis appears to match that of [12], affinity propagation with different parameters provided a more refined clustering, possibly capturing seven stages of myoblast differentiation (see also Supplementary Figure 3).

**Figure 4.**
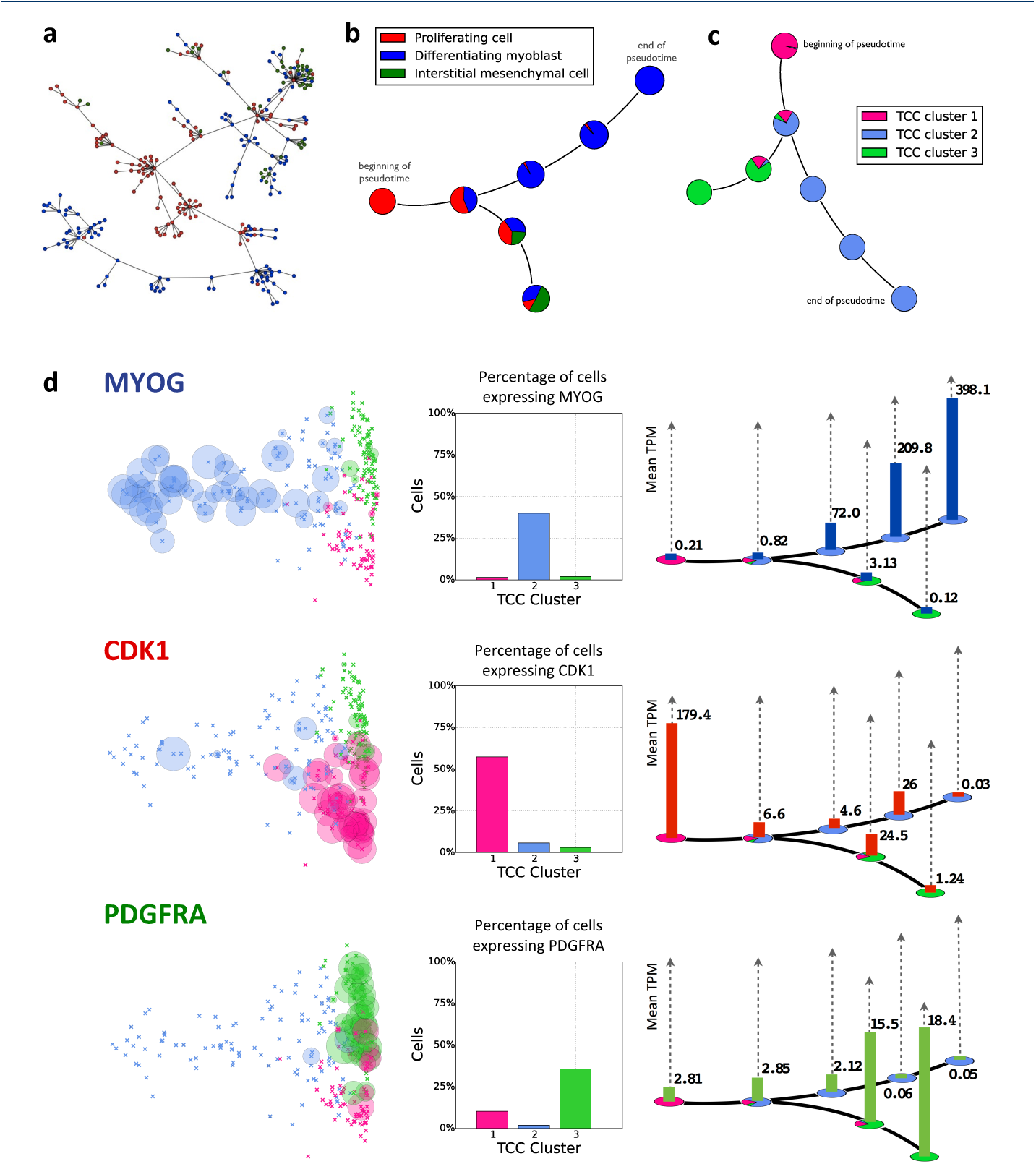
Temporal ordering of differentiating primary human myoblasts using transcript-compatibility counts. (a) A minimum-spanning tree (MST) was drawn through the 271 cells using Jensen-Shannon distances computed between 1,101, 805-dimensional vectors of TCCs of cells in the dataset. Following the longest path does not show a clear cell differentiation pattern. (b) Affinity propagation clustering generated 7 clusters (after collapsing spurious clusters with less than 5 cells into their nearest neighboring cluster), and an MST was drawn through the centroids of the clusters. Using the labels from Trapnell *et al*. [12], the longest path shows a differentiation pattern from proliferating cell (red) to differentiating myoblast (blue). The MST also shows how some proliferating cells alternatively differentiate into interstitial mesenchymal cells (green). (c) The cells were then clustered into 3 groups based on their transcript-compatibility counts, and the MST from b was re-labeled using these new cell types. (d) The expression levels of the genes *MYOG, CDK1*, and *PDGFRA* were analyzed for the 3 TCC clusters. *MYOG, CDK1*, and *PDGFRA* show greater expression for centroids from clusters with greater proportions of TCC cell types 1, 2, and 3, respectively. For each gene, a histogram over each centroid shows how expression level evolves with the differentiation process. *CDK1, MYOG*, and *PDGFRA* being markers for proliferating cells, differentiating myoblasts, and interstitial mesenchymal cells, indicate that the clustering and centroid-ordering based on TCC captures intermediate steps of the human myoblast differentiation trajectory.

A central idea in pseudo-temporal ordering of cells relies upon the construction of a minimum spanning tree (MST) over the pairwise distances of their corresponding gene expression vectors [48]. This attempts to capture the trajectory of a hypothetical cell that gradually “moves” through different cellular states or differentiation stages in a high-dimensional gene expression space. Our results show that the same concept can be applied to transcript-compatibility counts. A key step in Monocle is to *first* reduce the dimensionality of the data by independent component analysis (ICA) and *then* compute the MST based on Euclidean distances on the plane. Here we take a different approach and compute the MST on “cluster centers” in high dimensions (See Methods). Both approaches aim to battle the biological and technical noise that is inevitably introduced in scRNA-Seq experiments. Even though we could have used Monocle directly on transcript-compatibility counts, the design and comparison of specialized tools is beyond the scope of this paper.

Figure 4d validates the three primary clusters and the pseudo-temporal ordering obtained by our method based on three key myoblast differentiation markers, *MYOG, CDK1* and *PDGFRA* (see Supplementary Figure 4 for an additional set of genes taken from [12]). Interestingly, the expression of these genes gradually evolves over the pseudo-temporally ordered clusters, capturing both the underlying differentiation trajectory of proliferating cells to myoblasts, and the corresponding branching towards mesenchymal cells, as was observed in [12].

Finally, we should point out that although the three primary clusters of [12] are evident in our results, they are not identical. This naturally raises the question of whether clustering on (high-dimensional) transcript-compatibility counts could possibly lead to cell mis-classification. Our results show that this is not the case. In Figure 5 we investigated one cell that seemed to have been severely mis-classified by our method as a differentiating myoblast while it was identified as a proliferating cell by Monocle. However, an analysis of the expression levels of 12 marker genes obtained from [12] shows that this cell displays more similarity to differentiating myoblasts than proliferating cells. Overall our results seem to suggest that transcript-compatibility counts, being directly obtained from sequenced reads, might constitute a less noisy representation of the “transcriptomic state” of a cell compared to the one obtained by quantifying its gene expression.

**Figure 5.**
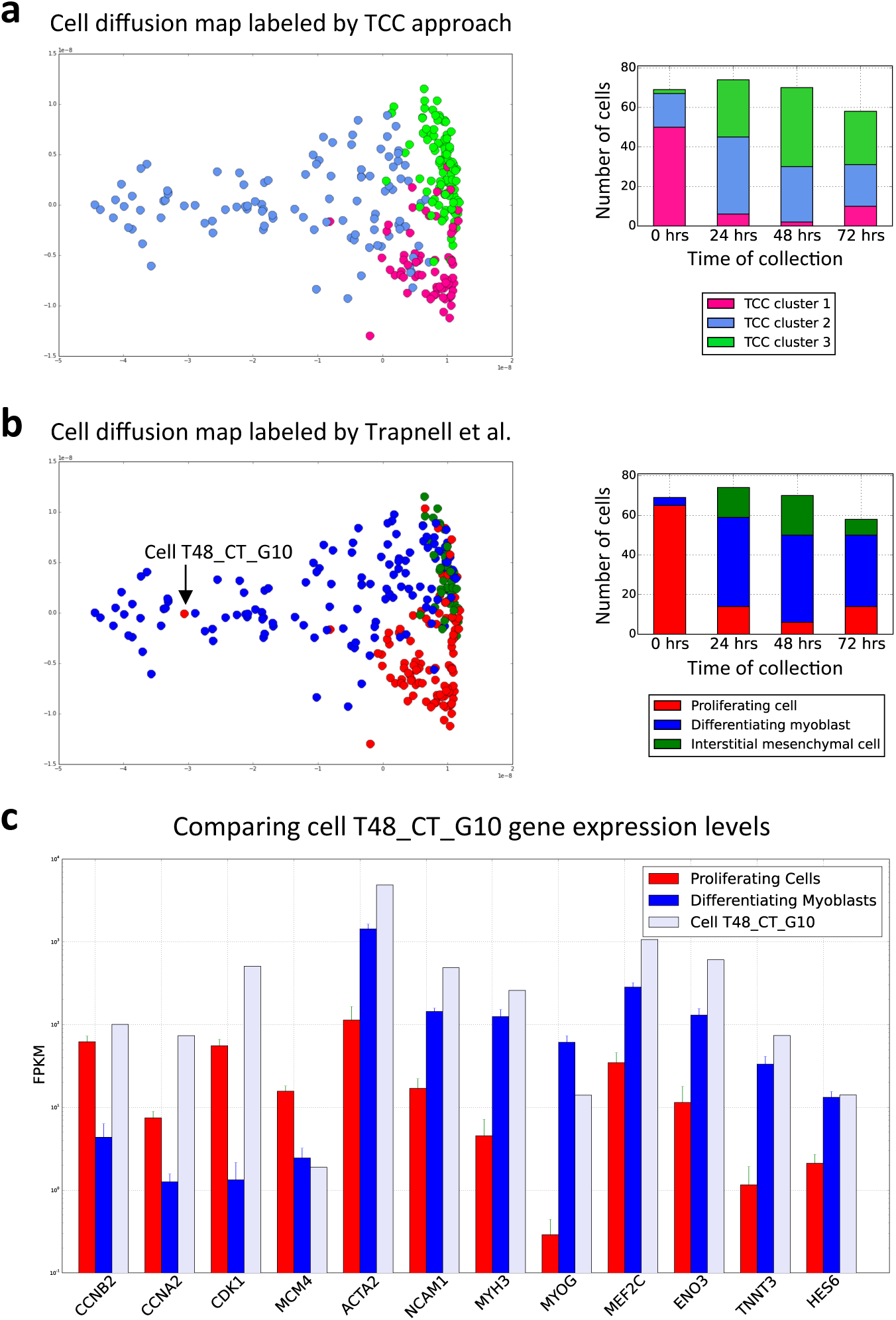
Clustering primary human myoblasts based on transcript-compatibility counts. (a) The transcript-compatibility counts matrix for 271 primary human myoblasts from [12] is visualized using a diffusion map. Three clusters obtained using affinity propagation are shown along with the distribution of these cells across the 4 cell-collection timepoints (0, 24, 48 and 72 hours), (b) The diffusion map obtained using transcript compatibility counts is relabeled using the cells reported by [12]. Clusters 1,2,3 generated by the transcript compatibility based method map to proliferating cells, differentiating myoblasts, and interstitial cells respectively. According to Trapnell *et al.’s* labels, the transcript compatibility based method seems to have severely misclassified cell T48_CT_G10 (SRR1033183) as a differentiating mycoblast. (c) Comparing the expressions of 12 differentiating genes in T48_CT_G10 with those of the average proliferating cell and the average differentiating myoblast, 8 out of the 12 genes show expressions similar to what one would expect from a differentiating myoblast. *MYOG* seems to show an FPKM of 14, which while more than the mean expression of proliferating cells (around 0.28) is much less than the mean expression of differentiating myoblasts (around 61.33). We note that this cell has the highest expression of *MYOG* among all cells labelled by Trapnell *et al*. as proliferating cell (and the second highest cell has expression around 5.4). However there are 88 differentiating myoblasts with MYOG expression less than 15 FPKM. Hence it is reasonable to think that this MYOG expression is more typical of differentiating myoblasts than proliferating cells. Only genes *CDKl* and *CCMB2* show expressions close to what one would expect from a proliferating cell. Even though *CDKl* is a highly specific marker for proliferating cells, the above gene profile indicates that classifying cell T48_CT_G10 as a differentiating myoblast seems reasonable.

### Cell classification in the mouse cortex and hippocampus

The re-analysis of [12] shows that clustering of transcript-compatibility counts can be useful on a single dataset, but we believe that the true power of our approach lies in its broad applicability to multiple single-cell assays. In contrast to the standard quantification pipeline, obtaining transcript-compatibility counts does not require a read-generation model; our method can be directly applied to a wide range of scRNA-Seq datasets and transcript-compatibility counts can be used to analyze sequenced reads without any assay-specific information. To make this point, we re-analyzed a recent large scRNA-Seq experiment published earlier this year [7] that uses an assay based on unique molecular identifiers (UMI). In contrast to [12] where paired-end reads were sampled from fragments covering the entire length of the transcripts, [7] used single-end reads that were only obtained from the 3’-end of the transcripts.

Zeisel et al. [7] examined a very diverse population of 3005 cells obtained from the cortical and hippocampal regions of the mouse brain. In order to analyze this complex dataset, the authors developed a state-of-the-art hierarchical bi-clustering method called BackSPIN (based on SPIN [30]) and were able to identify 47 distinct sub-populations of cells within nine major brain cell types. This fine-grained analysis also revealed a previously unknown post-mitotic oligodendrocyte sub-class, referred to as Oligo1 in [7].

Figure 6 shows the clusters obtained by applying our method to the above dataset and compares our method’s clustering accuracy to various quantification-based methods. In order to systematically assess the clustering accuracy, we iteratively sub-sampled cells from two different cell types at random and evaluated the ability of each method to distinguish between these types. Since the development of specialized clustering algorithms is orthogonal to our paper, we compared based on the same clustering algorithm throughout (see Methods). Our results indicate that transcript-compatibility counts can be more accurate than standard model-based RNA-Seq quantification tools (such as eXpress) that try to estimate the underlying read-generation model from the data. Our transcript-compatibility counts based method is in fact able to achieve similar accuracy with the assay-specific quantification approach used in [7] (that explicitly takes into account the significant 3’-end bias in this dataset). Clustering transcript abundance quantifications output by kallisto results in lower accuracy due to the mismatch between kallisto’s read-generation model and this dataset, further emphasizing the importance of using transcript-compatibility counts which are computed without using any such model.

**Figure 6.**
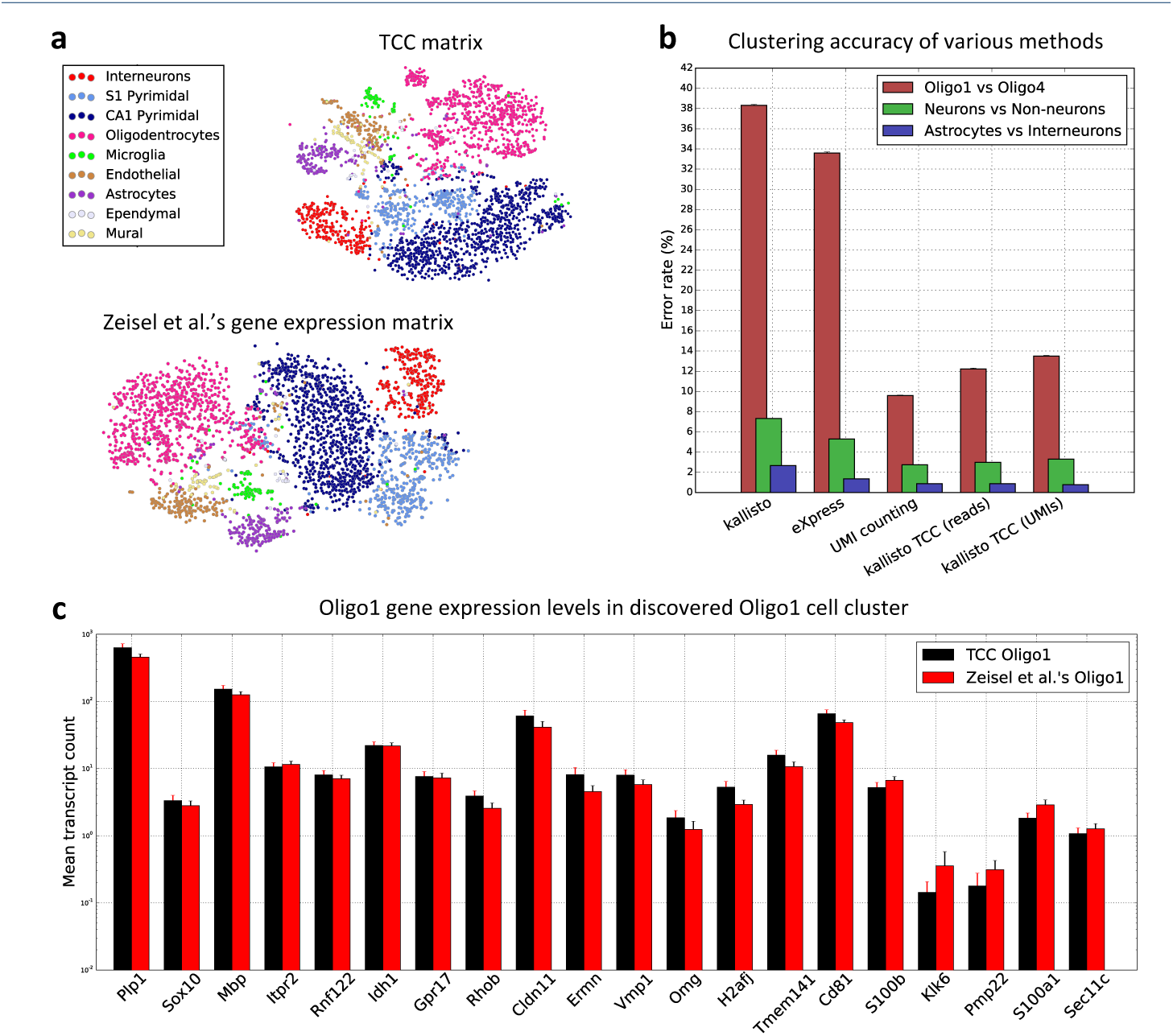
Clustering mouse brain cells based on transcript-compatibility counts. (a) The TCC distribution and gene expression matrices for the 3005 mouse brain cells are visualized using t-SNE (based on Jensen-Shannon distances between TCC distributions and gene expression distributions of cells respectively) and colored with the cell type determined by Zeisel *et al*. [7]. We note that the transcript-compatibility based t-SNE also visually maintains the cluster structure of the 9 major clusters, even though it can be computed two orders of magnitude faster than the gene expression matrix, (b) Cells from each of two cell types determined by Zeisel *et al*. were randomly selected, and then the clustering accuracy of multiple methods was tested. The clustering accuracy was measured as the error rate of the clustering. First, we note that the 3’-end bias in this dataset significantly affects the accuracy of kallisto and eXpress that have been chosen here as representative methods for model-based quantification (See Methods). For each point in the eXpress and kallisto curves, we took the minimum of the error rates obtained with bias modeling turned on and off. By avoiding estimation of the read model, transcript-compatibility based methods were indeed more accurate. We see that transcript-compatibility based clustering achieves similar accuracy to the gene-level UMI counting method implemented by the authors for this dataset without explicitly accounting for PCR biases. Refining transcript-compatibility counting to correct for PCR biases (by counting only the distinct UMI’s of reads in each equivalence class) leads to a marginal improvement of our method, (c) Running affinity propagation on the TCC distribution matrix (using negative Jensen-Shannon distance as similarity metric) produced a cluster of 28 cells, 24 of which were labelled Oligol. Zeisel *et al*. [7] classified 45 of the 3005 cells as this new class of cells. The bar plot compares the mean expression of selected oligodendrocyte marker genes in the TCC cluster to their mean expression in Zeisel *et al.’s* Oligol. As reported in [7], Oligol cells are characterized by their distinct expression of genes such as *Itpr2, Rnfl22, Idhl* and *Gprl7*. The similarity of the bars seems to suggest that clustering on TCC can capture this fine grained information. Note that although single cell clustering was entirely performed based on transcript-compatibility counts, the gene expression data used to evaluate this figure were obtained from Zeisel *et al*.

Quite remarkably, our method (via affinity propagation on all cells) was further able to recover the Oligo1 cluster of cells, showing that transcript-compatibility counts can indeed capture distinct cell signatures without actually quantifying their gene expression (Figure 6, Methods). Overall, in our experiments we observed that unsupervised clustering of transcript-compatibility counts typically yielded more than 47 clusters, which was also the case in [7]. Some of our clusters were very small, probably capturing outlier cells, while others seemed to be further splitting the 47 cell subtypes identified in [7].

To further investigate this, we focused on another oligodendrocyte sub-population, referred to as Oligo3 in [7]. As reported in [7], Oligo3 cells were almost exclusively observed in the somatosensory cortex and were identified by the authors as being in an intermediate stage of maturation – in between premyelinating and myelinating oligodendrocytes. Even though the Oligo3 cells appear to be well-clustered together, as visualized by t-SNE (Figure 7a), affinity propagation on transcript-compatibility counts with various parameters consistently separated them into two sub-clusters. Our results in Figure 7b seem to suggest that a sub-population of Oligo3 cells (captured by one of our sub-clusters) expresses an unusual signature of endothelial/vascular genes on top of the expected myelin related genes. Interestingly, similar findings have been reported recently in [37], suggesting a possible (experimental) contamination of several oligodendrocyte cells in the dataset at hand.

**Figure 7.**
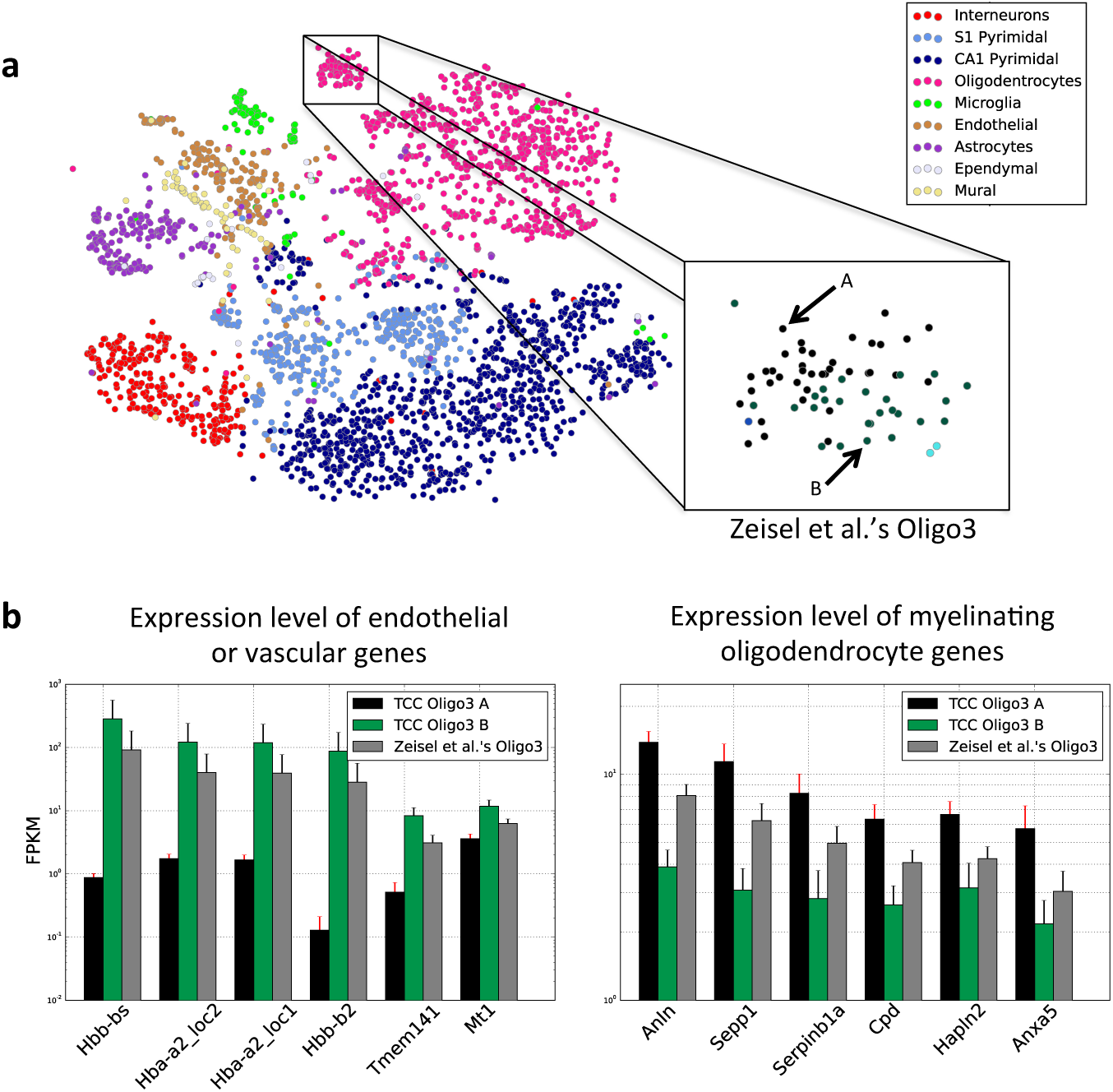
Analysis on transcript-compatibility counts refines the classification of mouse brain cells. (a) Runs of affinity propagation with different propagation and damping parameters were carried out on the TCC matrix for the 3005 mouse brain cells of [7]. The Oligo3 subclass discovered by Zeisel *et al*. was consistently split into to subclasses A and B. (b) Cells in the Oligo3 A class showed greater expression of endothelial/vascular genes and lower expression of myelinating oligodendrocyte genes. The opposite was true for the cells in Oligo3 B. This result may corroborate the potential contamination of oligodendrocytes in the Zeisel *et al*. dataset that has recently been reported in Fan *et al*. [37].

## Discussion

In this paper we introduced a novel method that uses transcript-compatibility counts – instead of gene expression profiles – as distinct cell signatures for clustering single cell data. Note, however, that the main focus of our method is not about how to cluster (i.e., the particular choice of clustering algorithms), but rather *what* to cluster on. To emphasize this point we considered simple, “off-the-shelf” clustering methods, that directly use the corresponding TCCs as their input. Interestingly, while these methods may not be able to recover accurate clusters when applied to gene expression vectors (see Supplementary Figure 6 or [7, Figure S3] for example), our results showed that TCCs maintain all the necessary information to recover the analyses of [12] and [7].

Even though clustering alone can reveal important information about a single-cell RNA-Seq experiment, further biological interpretation of the results (marker gene identification or differential expression) requires some form of quantification of expression profiles within and in between clusters. So it is natural for one to think that eventually the quantification bottleneck will still manifest itself in single-cell analysis. A key observation however is that given an accurate clustering of the cells, each and every individual cell’s gene expression profile is no longer needed; one can extract an accurate *statistical* representation of the gene expression within each cluster – without having to quantify all cells separately. In particular, one can quantify the aggregate gene expression in each cluster (cluster centers) by pooling single-cell TCCs together and further estimate the corresponding gene variability by subsampling and quantifying only a few cells per cluster. For example, in Supplementary Figure 5 we used kallisto to quantify subsampled cells and the corresponding cluster centers (after clustering on TCCs) for the Trapnell *et al*.’s dataset and generated results that are very similar to the ones obtained in Figure 4 (where the corresponding gene expression profiles were obtained from [12]). Our method can therefore be used to effectively *reverse* the quantification and clustering steps in the conventional pipeline and potentially provide further end-to-end processing gains, depending on the needs/goals of each scRNA-Seq experiment. Overall, we believe that clustering before quantifying is a promising future direction for scRNA-Seq analysis which may lead to more robust and accurate quantification algorithms.

## Conclusions

The extraordinary developments in single-cell RNA-Seq technology over the past few years have demonstrated that “single-cell resolution” is not just a gimmick but an unprecedented tool for probing transcriptomes that can reveal the inner-workings of developmental programs and their resulting tissues. However the computational challenges of scRNA-Seq analysis, already very high due to the large number of cells to analyze, have been further exacerbated by the smorgasbord of assays that each introduce unique technical challenges.

The new method we have proposed and evaluated in this paper, namely analysis of scRNA-Seq based on transcript-compatibility counts, offers an efficient, accurate and broadly applicable solution for extracting information from scRNA-Seq experiments. In the same way that single-cell analysis can be viewed as the ultimate resolution for transcriptomics, transcript-compatibility counts are the most direct way to “count” reads. While we have focused on clustering of cells in this paper, we believe that transcript-compatibility counts may have applications in many other sequencing-based assays, and that further development of methods based on such counts offers a fruitful avenue of exploration.

The ability to obtain transcript-compatibility counts by pseudoalignment is a benefit that has its own implications and applications. For example, the speed of pseudoalignment facilitated quick experimentation with our method, and in assessing our accuracy on different datasets one discovery was that much less sampling than is currently performed is necessary to cluster cells. In the re-analysis of [7], we found that the main results, namely the clustering of cells and identification of cell types, were achievable with only 1% of the data (see Figure 8a and Supplementary Figure 1). This observation has significant implications for scRNA-Seq as it suggests that for clustering of cells, low-coverage sequencing may be sufficient thus allowing for larger experiments with more cells. Moreover, this low-coverage clustering performance can be achieved using our method, which is not tailored to the specific scRNA-Seq assay.

**Figure 8.**
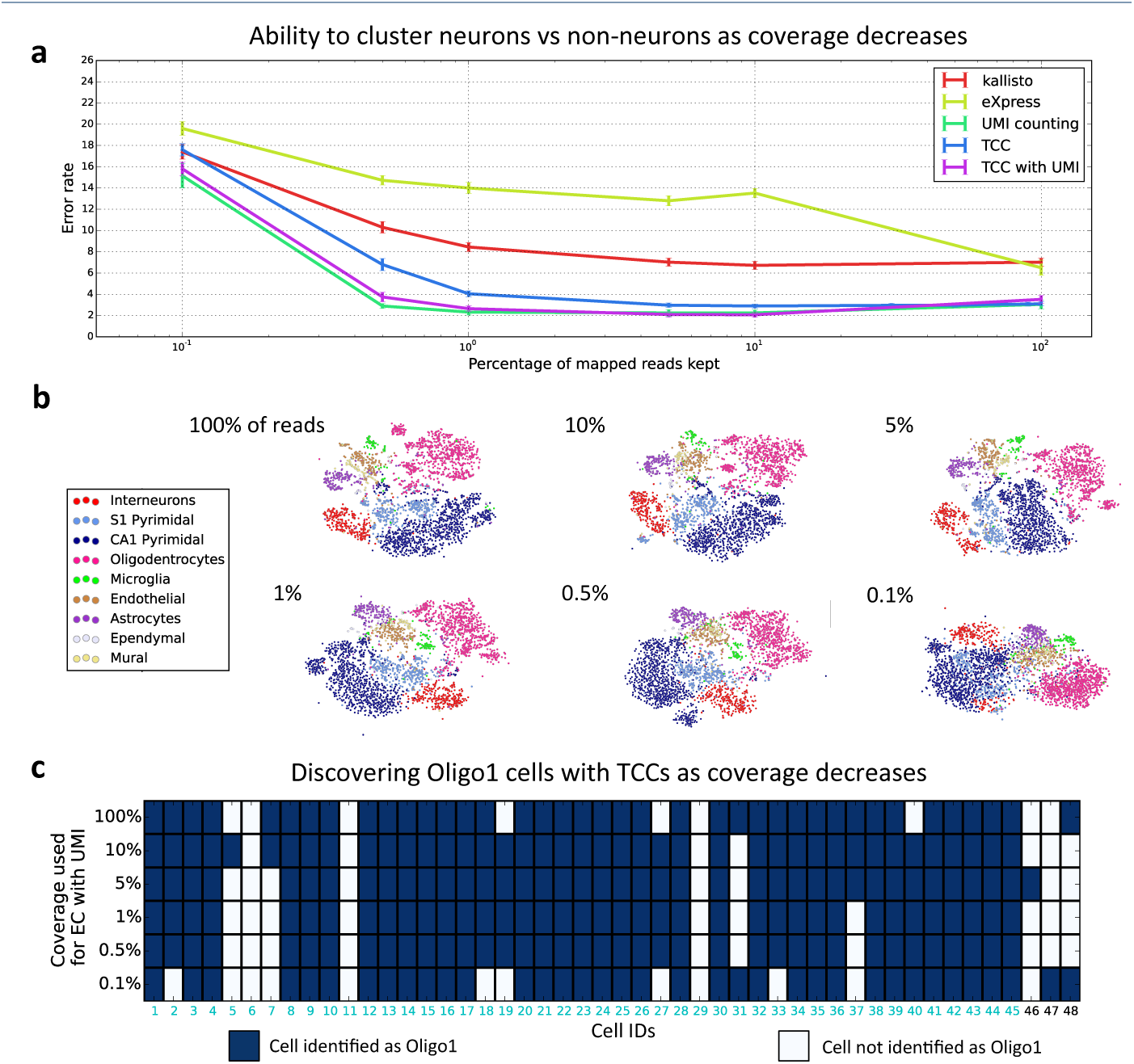
Coverage requirements for clustering based on transcript-compatibility counts. As an intermediate between raw reads and quantified transcript abundances, transcript-compatibility counts intuitively have more information than the reconstructed transcripts and less noise than the raw reads, (a) As the read coverage of a cell of the dataset of [7] decreases from approximately 627fc mapped reads, different methods have varying robustness to the loss of coverage. Each method was evaluated on its ability to cluster 200 randomly selected neurons mixed with 200 randomly selected non-neurons into the two cell types (the clustering of [7] being considered as the ground truth). Amongst methods which do not explicitly account for PCR bias, TCC based clustering performed much better than kallisto and eXpress and was quite close in performance to the UMI counting method of [7]. For each point in the eXpress and kallisto curves, we took the minimum of the error rates obtained with bias modeling turned on and off. By counting the number of unique UMIs rather than reads (TCC with UMI in the plot), transcript-compatibility based clustering was adapted to account for PCR bias, resulting in similar performance to that of gene-level UMI counting used in [7]. (b) Even at significantly decreased coverage depths, our method maintains clusters corresponding to the 9 major cell types identified by Zeisel *et al*. The transcript-compatibility disturbution matrices at varying coverage depths are visualized using t-SNE. (c) At various coverage depths, transcript-compatibility counting with UMIs disagrees slightly with the cells the authors labeled as Oligol (cyan cell IDs). As the coverage decreases, transcript-compatibility based affinity propagation still identifies a cluster that captures the vast majority of Oligol cells in the 3005-cell population.

## Methods

The code used to generate the results presented in this paper is available online on GitHub [49]. The Mus musculus transcriptome assembly used was GRCm38. The Homo sapiens transcriptome assembly used was GRCh38. The reference genome used for HISAT was build 10 of the mouse genome (mm10) from the UCSC genome browser.

### Computation of transcript-compatibility counts

In our implementation of the method, we use kallisto to compute transcript-compatibility counts via pseudoalignment (avoiding the quantification step that is usually performed when running kallisto altogether). In particular, we utilized the “pseudo” option of the kallisto RNA-Seq program which computes equivalence classes of reads after pseudoalignment. We used kallisto version 0.42.3 with k set to kallisto’s default value of 31. Even though kallisto pseudoalignment is a natural approach to obtain transcript-compatibility counts, one can in principle extract the same information from exact read alignments. To explore this alternative, we used HISAT (with the no-spliced-alignment option enabled) to align reads on the mouse transcriptome (GRCm38) in the case of Zeisel’s dataset and the human transcriptome (GRCh38) in the case of Trapnell’s dataset. Then, we generated the corresponding “alignment-based TCCs” by directly counting the number of multi-mapped reads aligned to each set of transcripts, and evaluated their performance in Supplementary Figure 8.

### Transcript-compatibility counts based on UMI information

The dataset of [7] has reads with unique molecular identifiers (UMIs). UMIs are typically used in scRNA-Seq to correct for PCR bias; biological copies of a transcript (distinct molecules) can be identified based on their UMIs. This information can be utilized in generating the transcript-compatibility counts from equivalence classes. Instead of counting all the reads in each equivalence class, we only count the reads with distinct UMIs. Transcript-compatibility counts with UMIs are shown in Figure 6b and Figure 8a (represented as “TCC with UMI” in the figures).

### Clustering Methodology

On obtaining the transcript-compatibility counts for each cell, we normalize by the total number of mapped reads to obtain a probability distribution called the transcript-compatibility count distribution or TCC distribution. We then compute the square-root of the Jensen-Shannon divergence [50] between the TCC distributions for each pair of cells. As a distance metric which satisfies the triangle [51] inequality, the square-root of Jensen-Shannon divergence is a natural choice for computing pairwise distances between two probability distributions. However, the results obtained here are not contingent on using the square root of Jensen-Shannon divergences as the measure of distances, and quite similar results are obtained when we use other distances between probability distributions such as the *ℓ_1_* distance to compute the pairwise distance matrix, (see Supplementary Figure 1). *ℓ_1_* distance (which is just twice the total-variation distance) in fact seems to perform better than Jensen-Shannon distance for low coverage (Supplementary Figure 1b). In contrast, Euclidean distance (*ℓ_2_* distance) seems to perform much worse (see Supplementary Figure 1). The fact that Euclidean distance is not a good distance metric to measure distances between probability distributions is widely documented (see for instance [52]).

All clustering carried out in this paper were done using off-the-shelf clustering methods.

We used spectral clustering using the pairwise distance matrices when we know the number of clusters in the data. This includes Figures 6b, 8a, and Supplementary Figure 1b with 2 clusters for the pairwise distance matrix (from TCC distributions) obtained for the data from [7].

The clustering method used when the number of clusters is not known is affinity propagation [53]. This is an unsupervised clustering algorithm based on message passing, which needs a pairwise similarity matrix as input. The pairwise similarity matrix is computed as the negative of the pairwise distance matrix that was computed.

To evaluate the clustering accuracy of our method in Figure 6b, we performed binary classification tests using the labels reported in [7] as the ground truth. In particular, we randomly sub-sampled two different types of cells and evaluated the ability of each pipeline to separate them into two clusters via spectral clustering. We performed these binary classification tests between 1) the sub-classes Oligo1 (45 cells) and Oligo4 (106 cells), 2) the cell types Astrocytes (198 cells) and Interneurons (290 cells), and 3) the more general cell types neurons (1628 cells) and non-neurons (1377 cells). The error rates for each test were obtained by randomly sampling 22, 99 and 200 cells from each of the two labels respectively, averaged over 10 monte-carlo iterations.

For clustering the dataset of [7], we used affinity propagation with the default parameters which set the preference value equal to the median of the similarity scores and the damping parameter equal to 0.5. On doing this, we obtained 89 clusters. Of the 89 clusters obtained, cluster number 22 had the largest match with the set of cells the authors labeled as Oligo1 (which was the new type of cells discovered in [7]). 24 out of the 28 cells in the cluster were labeled Oligo1 by [7]. There were a total of 45 cells labeled Oligo1 in [7] out of the total of 3005 considered. This is investigated in Figure 6*c*.

Also, affinity propagation with different parameters seems to split the class labelled Oligo3 in [7] into 2 classes. This is investigated in Figure 7, where the two classes considered were classes obtained with parameters set as before.

For clustering the dataset of [12], we used affinity propagation with preference parameter set to 1.3 and damping parameter set to 0.95 to obtain three clusters in Figure 4. To obtain 8 clusters on the dataset of [12], we used affinity propagation with preference parameter set to 0.6 and damping parameter set to 0.95 to obtain 8 clusters. Then after collapsing any cluster with less than 5 cells into the cluster closest to it, we obtain the seven clusters investigated in Figure 5. More details regarding our parameter choices for affinity propagation in this dataset are provided in Supplementary Figure 7.

### Partial order on clusters

On the [12] data set, for the seven clusters obtained, we first find the centroid TCC distribution of each cluster as the mean TCC distribution of all cells in the cluster. Then, we compute the pairwise Jensen-Shannon distances between the centroid TCC distributions (cluster centers). We then run a minimum weight spanning tree on the complete graph between the cluster centers with weights given by the computed pairwise distances. This gives us a partial-order on the clusters, which is investigated in Figure 4 and Supplementary Figure 3.

### Quantification

In this paper, we used kallisto and eXpress as representative methods for model-based quantification as they demonstrate similar accuracy to other available quantification tools [41]. In particular, for Zeisel et al.’s dataset (Figures 6b and 8a) we used these tools as a “negative control” to demonstrate the importance of the read-generating model when using quantified abundances to cluster single cell data. Although the significant mismatch between the assumed model (i.e., full transcript length coverage) and the 3’-end bias in this dataset is expected to affect the quantification accuracy for each individual cell, our results show that such a mismatch further impacts the accuracy of clustering and cell-type classification. To evaluate the cell-type classification performance of kallisto and eXpress, we took the minimum of the error rates obtained with bias modeling turned on and off.

For the Trapnell *et al*. dataset, we used kallisto to quantify transcript abundances and obtain the gene expression profiles within the clusters obtained from TCCs. Note that the read-generating model in this dataset is similar to the standard RNA-Seq model that kallisto uses for quantification. More specifically, in Supplementary Figure 5a we quantified the corresponding cluster centers by running kallisto’s EM algorithm on the pooled TCCs of each cluster. Using kallisto in this setting resembles bulk RNA-Seq quantification applied to the pooled reads coming from each individual cluster (instead of the entire population of cells). In Supplementary Figure 5b we further quantified randomly sub-sampled cells (20 cells per cluster) to obtain an accurate estimate of the gene expression variability within each cluster.

### Visualization of cells and clusters

We used t-SNE [54] to visualize the cells and clusters in Figures 6a, 8b, and Supplementary Figure 1a.

The left panel of Figure 4d, 5a, 5b and Supplementary Figure 4a was created using an implementation [55] of the diffusion map algorithm of [56].

## Competing interests

The authors declare that they have no competing interests.

## Author’s contributions

VN,GMK and JZ conceived the idea of clustering without quantification, performed analyses of data, analyzed and interpreted results and wrote the manuscript. DNT and LP interpreted results, supervised the project and wrote the manuscript.

## Acknowledgements

We thank Pall Melsted for implementing the pseudo command in kallisto. This is the command that allows for direct output of transcript-compatibility counts via pseudoalignment. Thanks to Bo Li for useful discussions about single-cell RNA-Seq assays and their biases. GMK and JZ are supported by the Center for Science of Information, an NSF Science and Technology Center, under grant agreement CCF-0939370. VN is supported in part by the Center for Science of Information and in part by a gift from Qualcomm Inc. LP is supported in part by the National Human Genome Research Institute of the National Institutes of Health under award number R01HG006129. DNT is supported in part by the Center of Science of Information and in part by the National Human Genome Research Institute of the National Institutes of Health under award number R01HG008164.

## Author details

^1^Department of Electrical Engineering and Computer Sciences, University of California, Berkeley,. ^2^Department of Electrical Engineering, Stanford University,. ^3^Departments of Mathematics and Molecular and Cell Biology, University of California, Berkeley,.

## Supplementary Figures for

“Fast and accurate single-cell RNA-Seq analysis by clustering of transcript-compatibility counts”

**Supplementary Figure 1:**
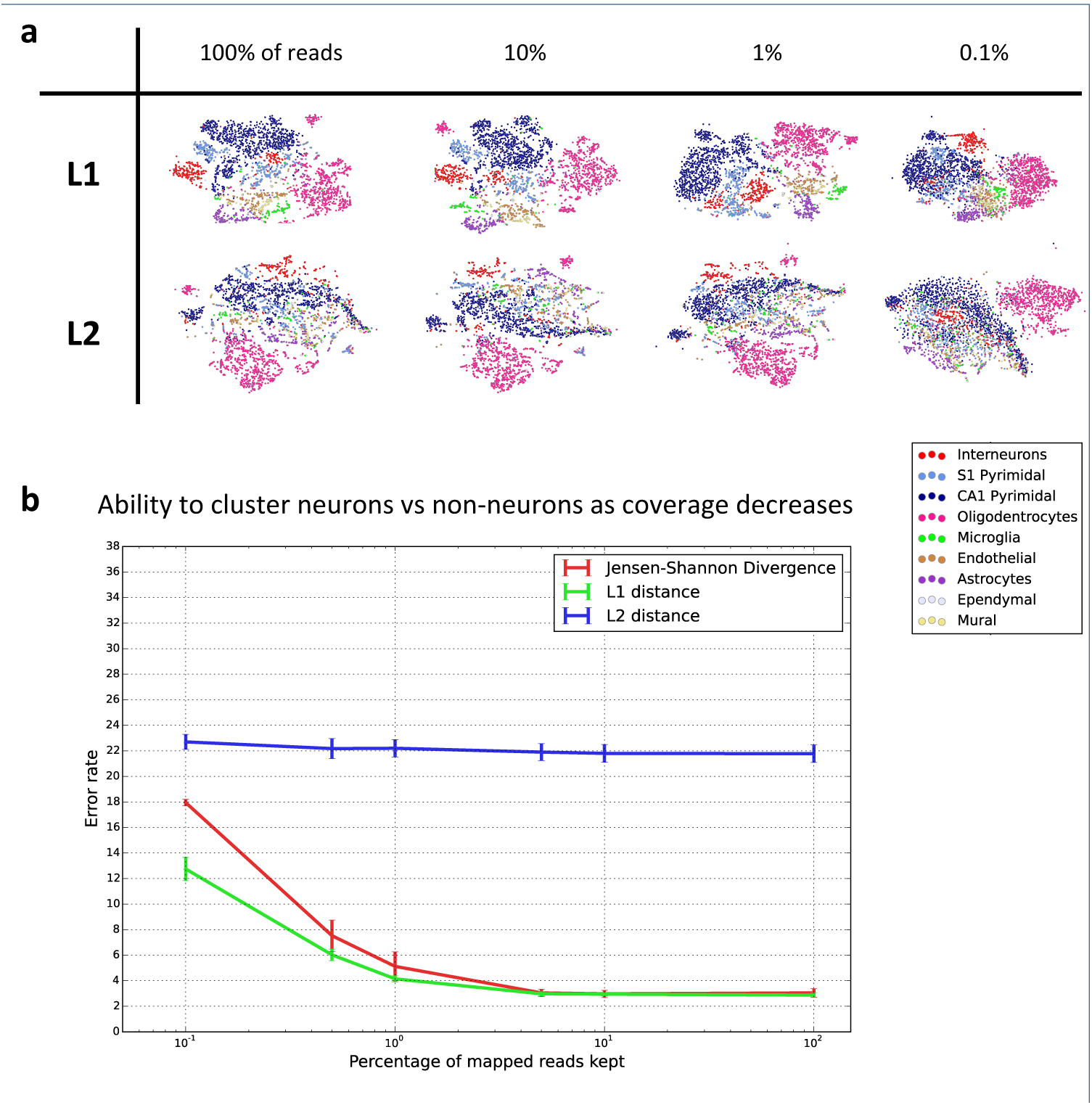
Comparison of different distance metrics to use to compute pairwise distances. (a) Shows the t-SNEs obtained when using *ℓ_1_* (Manhattan distance or twice the total variation distance) and *ℓ_2_* distances (Euclidean distance) instead of Jensen-Shannon distances to compute pairwise distances between TCC histograms obtained for 3005 mouse brain cells of Zeisel *et al*. [1]. The *ℓ_1_* distance seems to maintain the cluster centers to a much larger extent than *£2* distances, (b) As the average read coverage of each cell in the dataset decreases from approximately 627,000 mapped reads, spectral clusterings based on different distance metrics exhibit varying ability to distinguish neurons from non-neurons. While both *ℓ_1_* distance and Jensen-Shannon Divergence perform similarly well at high coverage (error rate 5%), the commonly used *ℓ_2_* distance resulted in significantly worse performance. We note that *ℓ_2_* distance is known to be a bad metric to use while comparing probability distributions. For the two classes picked, *ℓ_1_* distances perform better that Jensen-Shannon distance at low coverage.

**Supplementary Figure 2:**
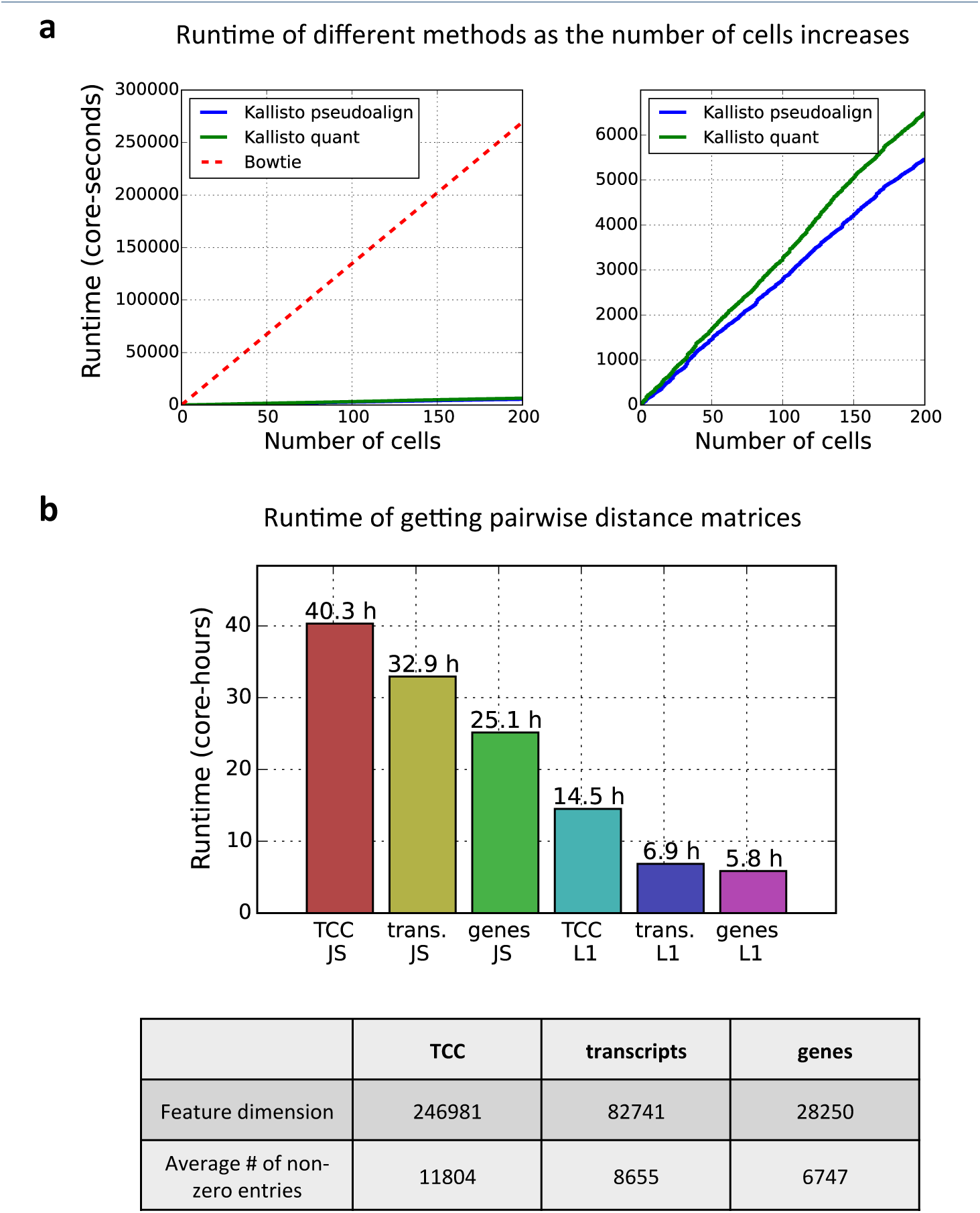
Runtimes of bowtie, kallisto quant, kallisto-pseudoalign, and computation of pairwise distance matrices. (a) The runtimes of Bowtie, kallisto with both pseudoalignment and quantification (kallisto-quant), and kallisto with just pseudoalignment (kallisto-pseudoalign) were obtained for 200 randomly selected cells from Zeisel *et al.’s* 3005 mouse brain cell dataset [1] as shown on the left pane. The (extrapolated) runtime of Bowtie was higher than the runtimes of the two pseudoalignment-based methods. When comparing kallisto-quant against kallisto-pseudoalign (as shown on the right pane), kallisto-pseudoalign is slightly faster, saving approximately 5 seconds per cell. As the number of cells scales up to 44, 000 for novel sequencing technologies such as DropSeq, kallisto-pseudoalign will have savings of about 60 hours compared to kallisto-quant and 1.8 years compared to bowtie. (b) The runtimes obtained for running pairwise distances on the distributions obtained from TCCs, transcriptome expressions, and gene counting are shown here. These times are shown for Jensen-Shannon distance and *ℓ_1_* distances. The feature dimension indicated in the table equals the number of features (either TCC, transcript abundances, or gene abundances) that are non-zero in at least one of the 3005 samples.

**Supplementary Figure 3:**
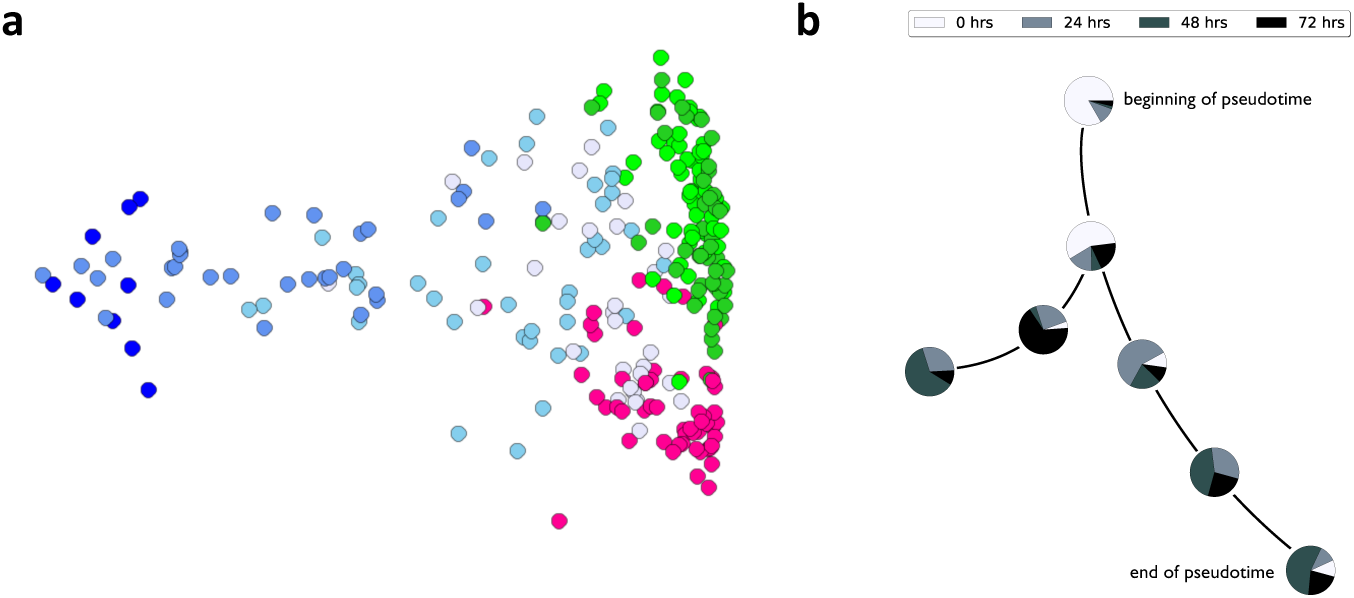
More details on the 7 clusters obtained from affinity clustering in the data-set of [2]. (a) Shows the diffusion map of cells colored by the labels of the 7 clusters. (b) Each pie-chart node in the MST shows the distribution of the cells of each cluster in real-time. The tree on which these are placed corresponds to the pseudotime obtained.

**Supplementary Figure 4:**
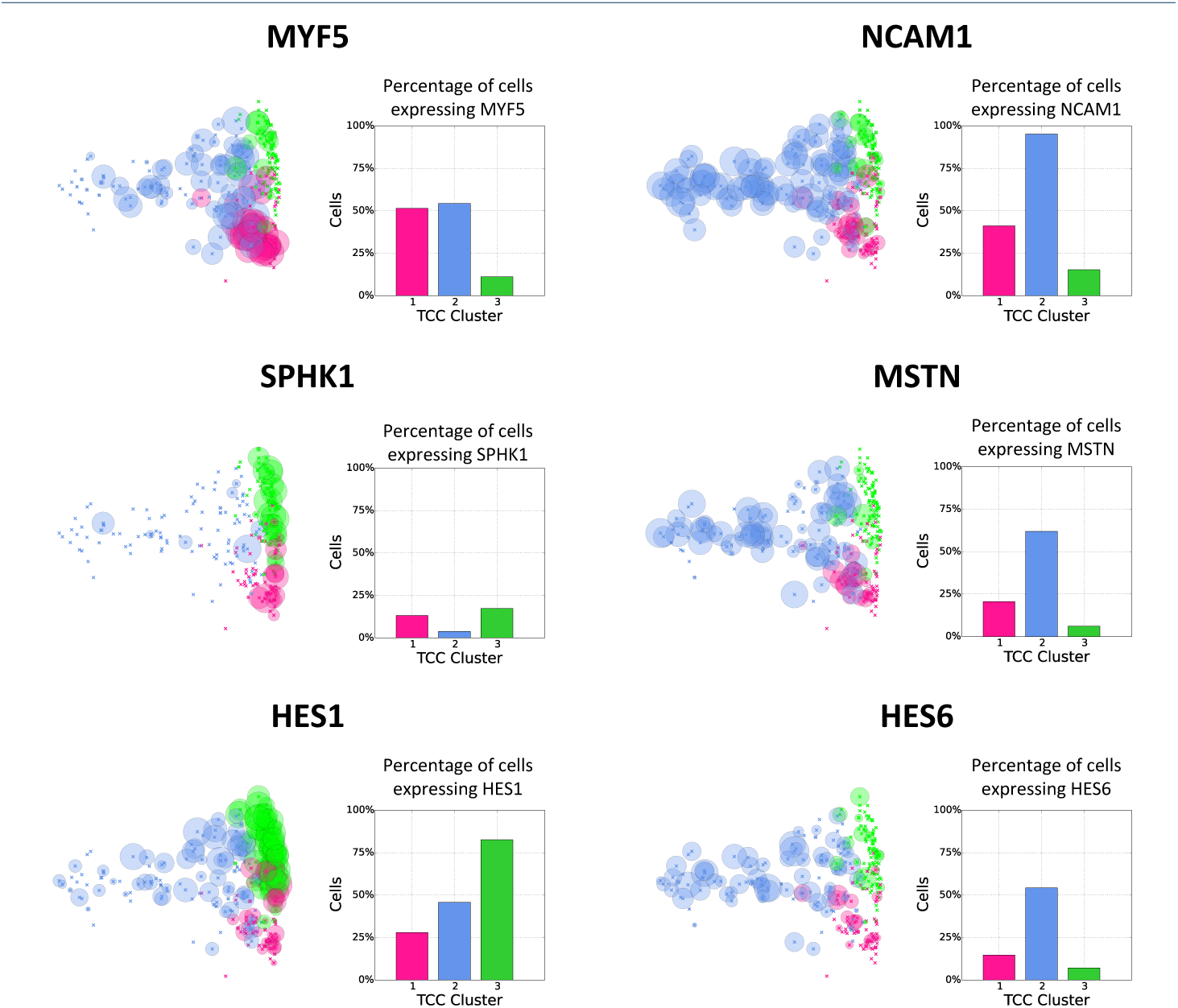
More genes to validate the 3 clusters obtained from the data-set of [2]. Shows the distribution of various other genes that are known to be markers of the three states represented by the three TCC clusters. The patterns discovered here using TCC closely matched those found by Trapnell *et al*.

**Supplementary Figure 5:**
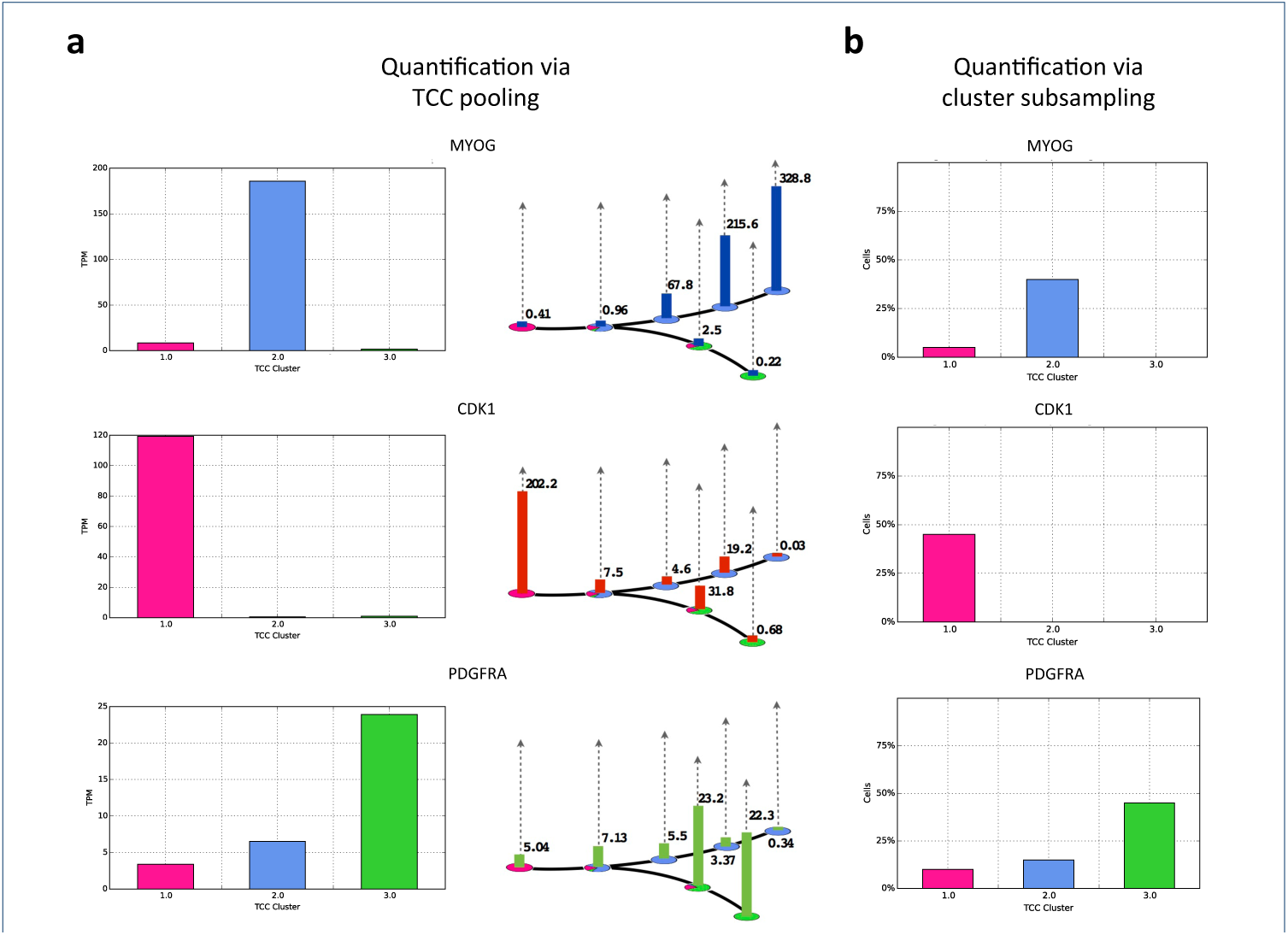
Quantifying after clustering to validate clusters obtained. (a) The expression levels obtained after running kallisto’s EM algorithm on the pooled TCCs of each cluster. The left pane shows the mean TPMs of the 3 clusters. The right pane shows the mean TPMs of the 7 clusters overlaid on the MST from Figure 4. We note that these were obtained by running the EM algorithm 3 times and 7 times, respectively (once for each cluster). We also note that the TPMs are similar to those of Figure 4d. (b) Here we show an estimate of the number of cells expressing the each of 3 genes. The expression levels are obtained by randomly sampling 20 cells from each cluster and quantifying them. We note that the numbers obtained are similar to those of the middle pane Figure 4d.

**Supplementary Figure 6:**
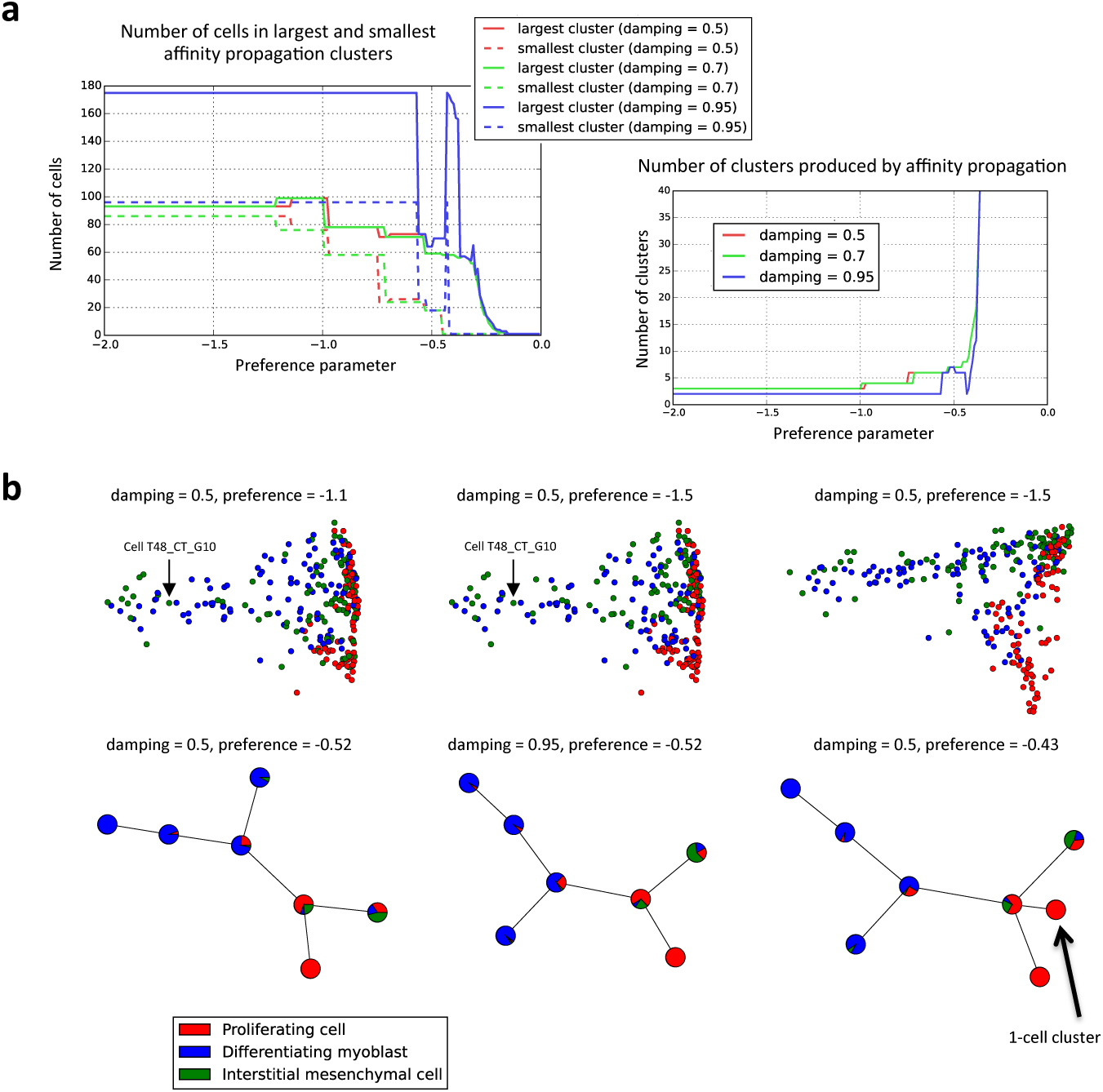
Selecting parameters for affinity propagation on [2]’s gene expression vectors. We note that choosing optimal parameters for affinity propagation requires some biological intuition, (a) For each of 3 damping parameter values, we swept through multiple preference parameter values. We looked for a combination of parameters that produced a reasonable amount of clusters roughly the same size. The left plot show two curves for each damping parameter: a dotted one indicating the number of cells in the smallest cluster and a solid one indicating the number of cells in the largest cluster. In the case where we do not know the correct number of clusters, we would use clusterings immediately before the large spike in number of clusters (right plot), resulting in about 7 clusters. The plots shown here are generated using Trapnell et al.’s expression vectors. We also noticed that from empirical testing, varying parameters in a flat region of the plot resulted in the exact same clusters, (b) There are multiple combinations of parameters that could generate 3 or 7 clusters. Here we use Trapnell et al.’s expression vectors to generate two MSTs. Each MST uses one of two combinations of damping and preference parameters selected based on the plots in (a). Slight tweaking of the preference parameters can result in an MST with 8 clusters, as shown in the right-most tree. Like we did in Figure 4, we would collapse the 1-cell cluster into its nearest cluster. Knowing that 3 cell types exist in the population, we also tried another two combinations of parameters to produce 3 clusters. For easy comparison to the TCC results in the main text, we visualized the clusters with the diffusion maps of Figure 4d. We see that the cell discussed in Figure 5 (T48_CT_G10) still fails to be classified as a differentiating myoblast. For additional comparison, we computed another diffusion map using Trapnell et al.’s expression vectors (right-most diffusion map).

**Supplementary Figure 7:**
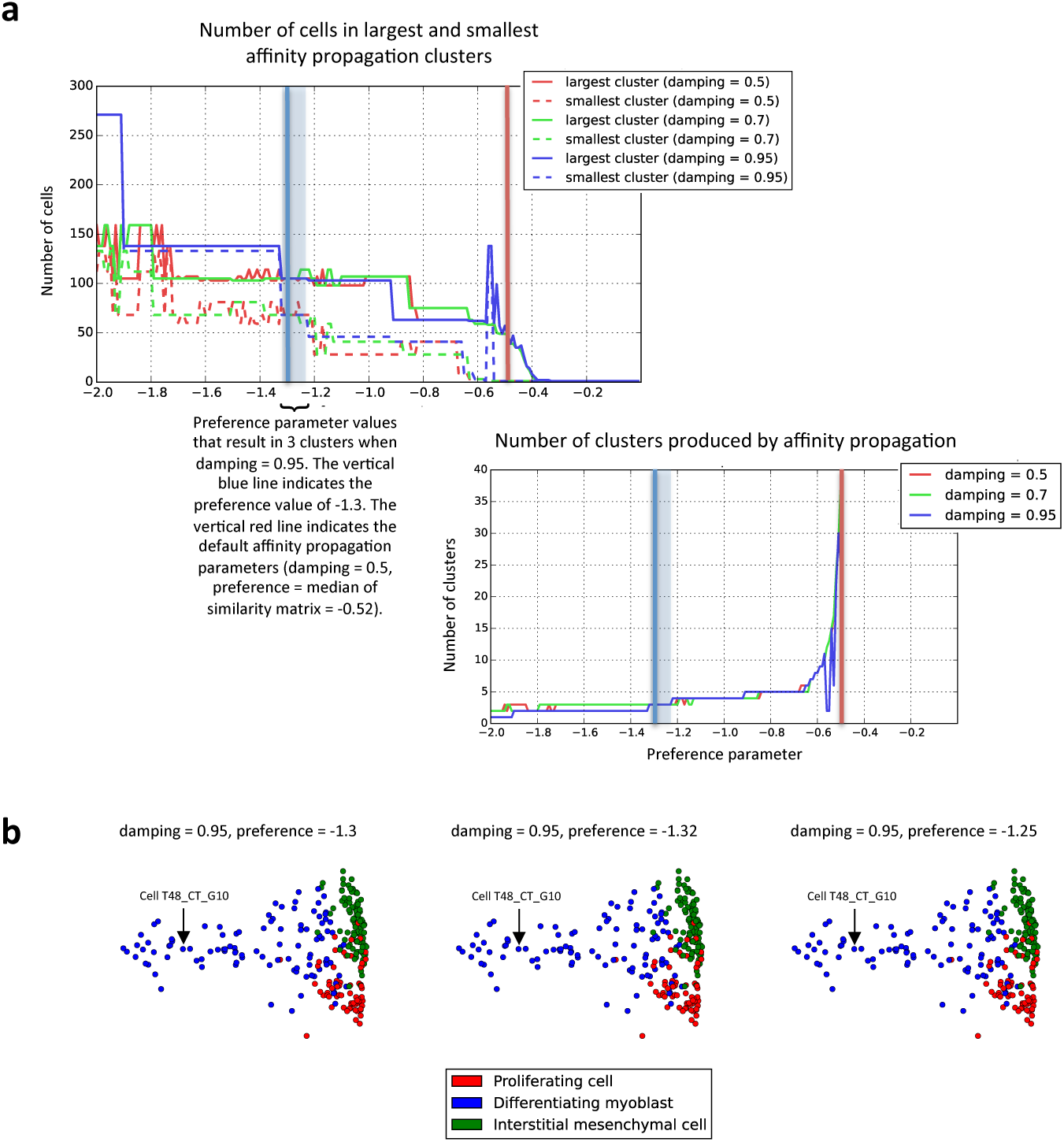
Selecting parameters for affinity propagation on TCC vectors for Trapnell’s dataset. (a) For the TCC approach, we performed the same parameter sweep presented in Supplementary Figure 6, resulting in the plots shown here. Additionally, we highlight the area of the curves where affinity propagation with a damping value of 0.95 results in 3 clusters. The default affinity propagation parameters of 0.5 for damping and -0.52, the median of the similarity matrix, for preference results in 24 clusters, 12 of which have only 1 member, (b) To test the stability of the clusters across a flat region of the curves, we looked at 3 combinations of parameters that resulted in 3 clusters when the damping value equals 0.95. The clusterings are identical, and we see that Cell T48_CT_G10 is consistently classified as a differentiating myoblast.

**Supplementary Figure 8:**
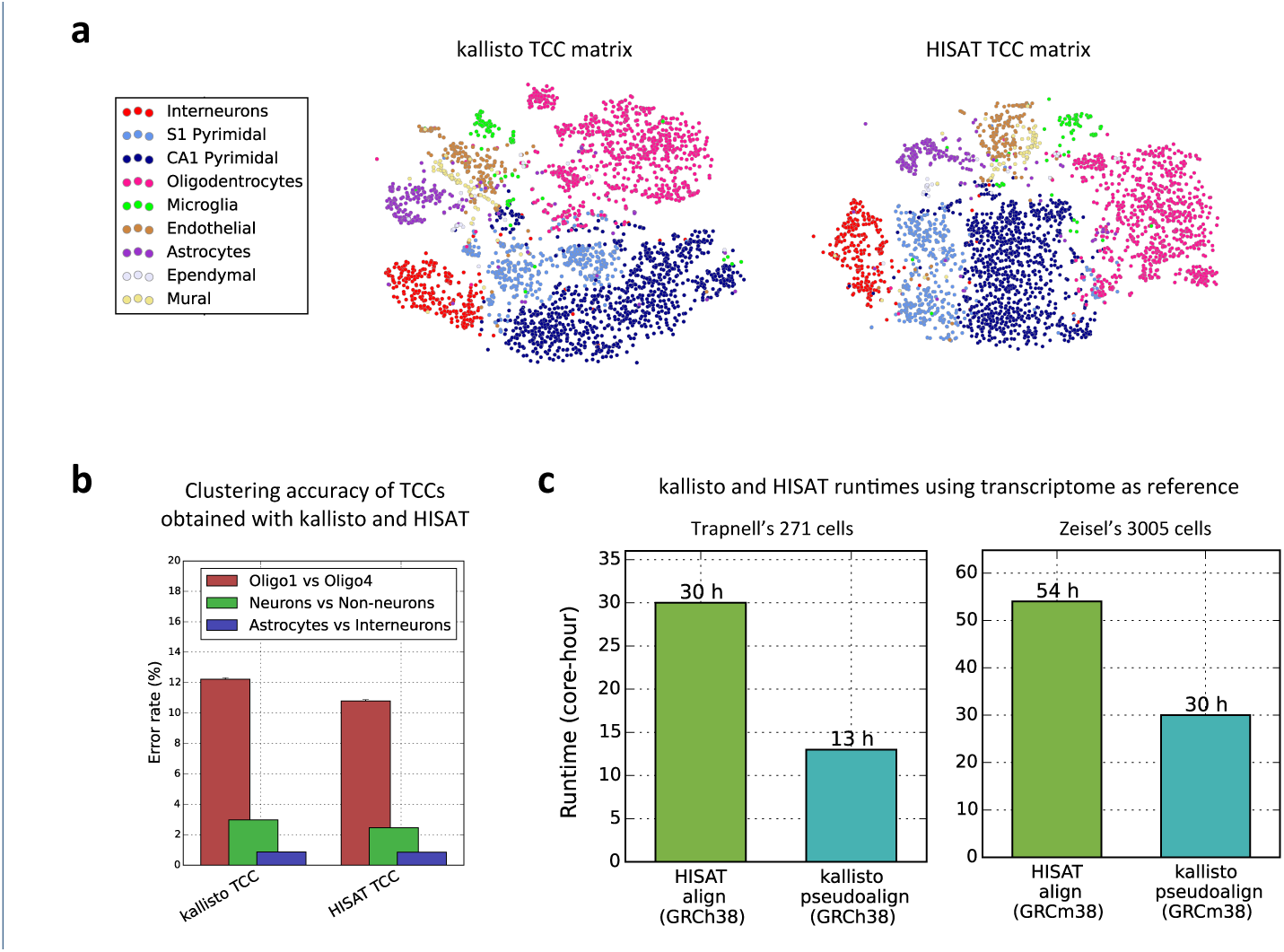
Comparing alignment-based TCC with pseudoalignment-based TCC. Alignment was performed using HISAT on the mouse transcriptome (GRCm38) in the case of Zeisel’s dataset and the human transcriptome (GRCh38) in the case of Trapnell’s dataset. HISAT’s --no-spliced-alignment option was used. TCC vectors can be generated from aligned reads by simply counting the number of ambiguous reads aligned to each set of transcripts. For Zeisel et al.’s dataset, HISAT maps 1, 843, 467,887 reads to 417, 515 equivalence classes, and kallisto maps 1, 768, 321, 229 reads to 246, 981 equivalence classes. We compare the (a) t-SNE visualizations on Zeisel et al.’s dataset, (b) clustering accuracies on Zeisel et al.’s dataset, and (c) runtimes of the two approaches on both Zeisel and Trapnell et al.’s datasets. Overall, alignment-based TCCs yield slightly better cell-type classification error rates on the Zeisel et al.’s dataset – at the cost however of a higher computation time.

